# Population Genomics of Nontuberculous Mycobacteria Recovered from United States Cystic Fibrosis Patients

**DOI:** 10.1101/663559

**Authors:** Nabeeh A. Hasan, Rebecca M. Davidson, L. Elaine Epperson, Sara M. Kammlade, Rachael R. Rodger, Adrah R. Levin, Alyssa Sherwood, Scott D. Sagel, Stacey L. Martiniano, Charles L. Daley, Max Salfinger, Jerry A. Nick, Michael Strong

## Abstract

Nontuberculous mycobacteria (NTM) pose a threat to individuals with cystic fibrosis (CF) due to an increased prevalence of pulmonary infections, innate drug resistance of the bacteria, and potential transmission between CF patients. To explore the genetic diversity of NTM isolated from CF patients within the United States (US) and to identify potential transmission events, we sequenced and analyzed the genomes of 341 NTM isolates from 191 CF patients as part of a nationwide surveillance study. The most abundant species in the isolate cohort were *Mycobacterium abscessus* (59.5%), followed by species in the *Mycobacterium avium* complex (37.5%). Phylogenomic analyses of the three *M. abscessus* subspecies revealed that more than half of CF patients had isolates in one of four dominant clones, including two dominant clones of *M. abscessus* subspecies *abscessus* and two dominant clones of *M. abscessus* subspecies *massiliense. M. avium* isolates from US CF patients, however, do not have dominant clones and are phylogenetically diverse. Longitudinal NTM isolates were compared to determine genome-wide single nucleotide polymorphisms (SNPs) that occur within patients over time. This information was used to compare between and within-patient SNP distributions, to quantitatively define SNP thresholds suggestive of transmission, and calculate a posterior probability of recent transmission given the SNP distance between two isolates from different patients. Out of 114 patients with *M. abscessus* subspecies, ten clusters of highly similar isolates from 26 patients were identified. Among the 26 patients in the *M. abscessus* clusters, 12 attended the same CF care centers. No highly similar isolate clusters were observed in *M. avium*. Our study reveals the contrasting genomic diversity and epidemiology of two major NTM taxa and the potential for between-patient exposure and cross-transmission of these emerging pathogens.

## Introduction

Nontuberculous mycobacteria (NTM) are ubiquitous environmental microorganisms found in water and soil that can cause severe human disease in susceptible individuals, including those who are immunocompromised ^1,2^ or have preexisting lung diseases such as cystic fibrosis CF ^3^. The predominant presentation of NTM disease is pulmonary infection, though extrapulmonary infections also occur ^3^. NTM infections are difficult to diagnose due to the slow growing nature of NTM and the requirement for specialized culture methods ^4^. NTM disease is difficult to treat because of the bacteria’s innate resistance to many antibiotics. Treatment courses can typically last months or years ^3^ with the potential for nonresponsiveness and adverse drug reactions ^5^.

NTM comprise over 190 mycobacterial species and subspecies ^6^, but only a limited number are clinically significant. US-based surveys find that the slowly growing species in the *Mycobacterium avium* complex (MAC; *i.e. M. avium, Mycobacterium chimaera*, and *Mycobacterium intracellulare*) and the rapidly growing species *Mycobacterium abscessus* (including its three subspecies; *M. abscessus* subsp. *abscessus, M. abscessus* subsp. *boletii*, and *M. abscessus* subsp. *massiliense*) are the most clinically relevant taxa associated with pulmonary infections ^7–9^. International studies suggest that NTM species vary geographically between countries and continents ^10^. A 2003 prospective US study of 986 patients from 21 CF treatment centers found that MAC species were the most prevalent NTM in CF patients with positive NTM sputum cultures (72% of patients), while 16% of patients had *M. abscessus* ^11^. A study of US CF Patient Registry data from 2010-2014 found that 61% of CF patients with positive NTM cultures had MAC species, primarily *M. avium*, and 39% had *M. abscessus* ^8^. The most recent CF Patient Registry Report from 2017 shows prevalence estimates of 51% for MAC species and 41% and for *M. abscessus* subspecies for CF patients who have had one or more positive NTM culture, suggesting that *M. abscessus* has significantly increased in prevalence in the CF community in the past 20 years ^12^.

Though MAC species are the most frequently observed NTM in US CF patients, information about the genetic diversity of clinical isolates lags behind that of *M. abscessus* ^13–17^. Much of the current understanding of *M. avium* diversity in the US is based on genotyping methods such as variable nucleotide tandem repeats VNTR ^18,19^ and serotyping ^20–23^, that lack the resolution of whole genome sequencing (WGS). Furthermore, genomic studies of *M. avium* are limited to non-CF isolate populations from Japan ^24,25^ and Germany ^26–28^. US-based genomic studies of *M. avium* and other MAC species recovered from CF patients are needed to better understand modes of acquisition and the impact of genotype on clinical outcome.

NTM were initially thought to be exclusively acquired from the environment until recent studies suggested that *M. abscessus* subsp. *massiliense* may be transmitted person-to-person ^13,29^. Intriguingly, this potentially “transmissible clone” of *M. abscessus* subsp. *massiliense* was also responsible for an epidemic of soft tissue infections in Brazil ^16,17^. Bryant *et al*. provided the first evidence, with genomic and epidemiologic data, suggesting patient-to-patient transmission of *M. abscessus* subsp. *massiliense* in a single CF center ^13^ and observed the presence of two dominant circulating clones of *M. abscessus* subsp. *abscessus* and one dominant circulating clone of *M. abscessus* subsp. *massiliense* throughout the global CF-NTM isolate population ^14^. Subsequent genomic studies from other European CF centers also found the same highly similar clones of *M. abscessus* subsp. *abscessus* and *M. abscessus* subsp. *massiliense* among different patients, but did not find epidemiological evidence of person-to-person transmission ^30,31^. This discrepancy, suggested that further studies were needed to determine the acquisition sources and transmission of such NTM isolates, especially in the US. It is also not known whether MAC species have dominant clones in the CF population and/or the potential to be transmitted person-to-person. To that end, we performed whole genome sequencing of NTM isolates sent from CF centers across the US. NTM isolates were received by the Colorado CF Research & Development Program (CF-RDP) at National Jewish Health in order to perform phylogenomic analysis, determine the population structure of NTM species affecting the CF population, and identify genetic evidence of potential transmission. Here, we provide an empirical investigation of 341 NTM isolates recovered from CF patients across the United States.

## Results

### Distribution of NTM species in the US CF population

The Colorado CF-RDP NTM isolate cohort included 341 isolates from 191 CF patients, representing 13 mycobacterial species (**Figure 1A, Table S1**). The three subspecies of *M. abscessus* made up 59.5% of total isolates, three species of MAC made up 37.5%, and the remainder included less frequently isolated NTM species with two or fewer isolates (2.9%). The taxa with the highest proportions of isolates *M. abscessus* subsp. *abscessus* (45.5%) followed by *M. avium* (23.5%). The high relative abundance of *M. abscessus* subsp. *abscessus* and *M. abscessus* subsp. *massiliense* in our study reflects the growing concern from CF centers about transmission of these taxa, and does not reflect the actual prevalence of these bacteria in the US CF population. ^8,11^. All *M. avium* isolates in the study were identified as *M. avium* subsp. *hominissuis* (distinct from *M. avium* subsp. *avium, M. avium* subsp. *silvaticum* and *M. avium* subsp. *paratuberculosis). M. intracellulare* was the second most abundant MAC species (10%), and *M. chimaera* was relatively rare (4%). At the patient level, a majority of patients had only one isolate (133/191 = 69.6%), 20 patients had two isolates (10.5%) and 38 patients had three or more isolates (19.9%) spanning a range of 4 to 1,137 days between the first and last isolate collection (**Figure 1B**). Of the 30 patients with three or more longitudinal isolates spanning at least 60 days, 10/30 (33%) had more than one NTM species over time (range **63** to 1,137 days). The distribution of the Colorado CF-RDP isolate cohort included patient isolates from 45 CF care centers in 22 states across the US, with geographic representation coast-to-coast and in northern and southern regions. Two-thirds of patients (124/191 = 64.9%) had samples sent from CF centers outside of Colorado, four patients has samples sent from CF facilities both in Colorado and out of state (4/191 = 2.1%), and approximately one third (67/191 = 35%) were referral patients to The Colorado CF Center (National Jewish Health/St. Joseph Hospital). National Jewish Health serves as a national referral center for mycobacterial infection, and many patients visit from out of state for diagnosis and treatment. As such, we assigned samples received from Colorado facilities with the actual patient state of residence, to more accurately investigate potential geographic influence on strain phylogeny and genomic relatedness (**Figure 1C**). Of the 71 patients treated at Colorado CF centers, 24 (33.8%) had a state of residence other than Colorado.

**Figure 1.**
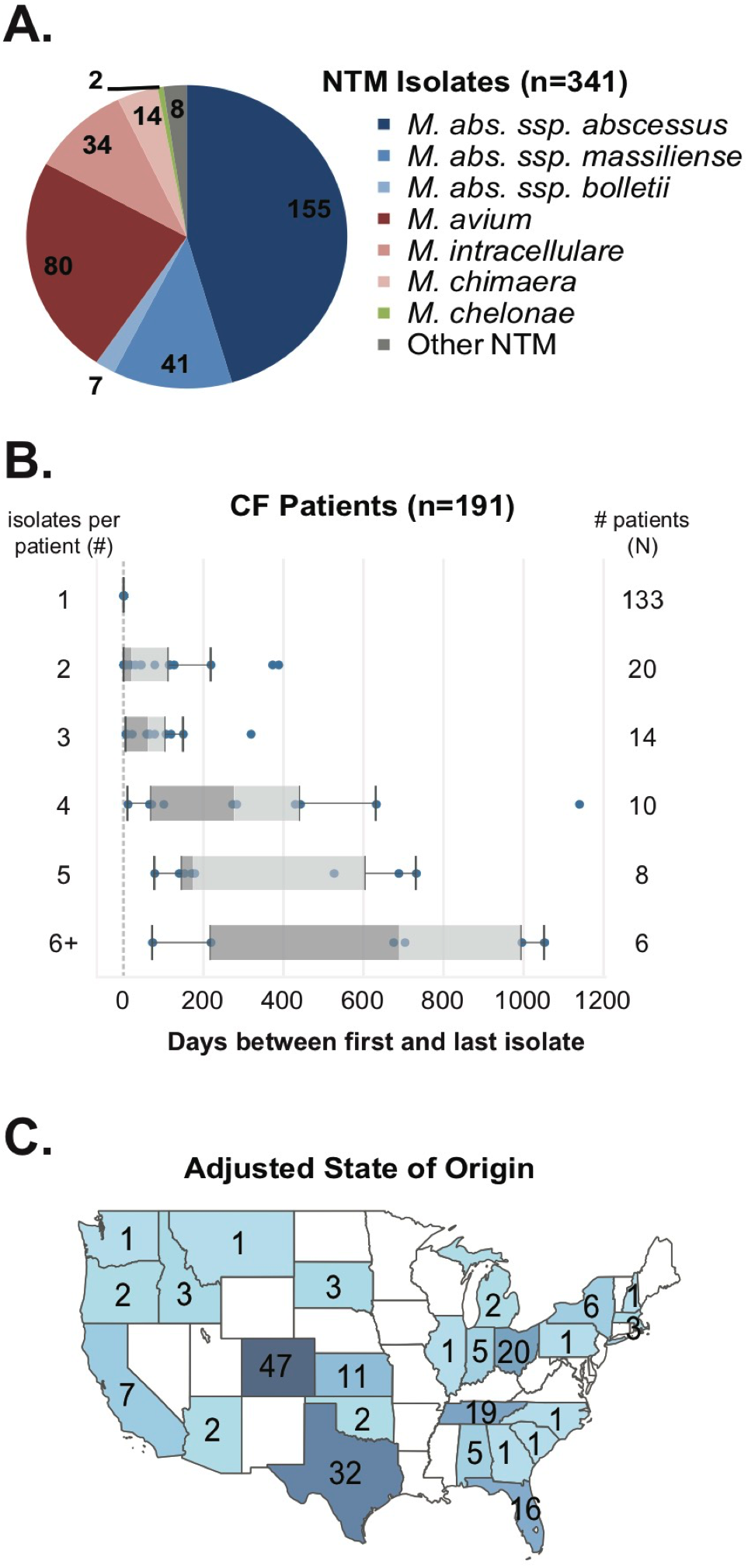
NTM isolate cohort of the Colorado CF-RDP with a total of 341 isolates from 191 patients and 13 different species. A. Distribution of NTM isolates by species. The most abundant species in the cohort belong to *M. abscessus* subspecies (blue colors) and *M. avium* complex (red colors). B. Numbers of isolates per patient in the Colorado CF-RDP NTM isolate cohort. Approximately 133/191 (70%) of patients are represented by a single isolate and 58/191 (30%) of patients have 2 or more longitudinally sampled isolates. C. Geographic distribution of patient states of origin (adjusted for known state of residence).

### Phylogenomic nomenclature to categorize relationships between isolates

To assign genetic relationships within each species/subspecies, we adopted the following hierarchical categories that represent the *least* to *most* genetic similarity: lineage, clone and cluster. ***Lineage*** denotes a monophyletic clade of independent isolates within a species ^24^. ***Clone*** denotes a dense monophyletic clade of isolates from different patients, different locations and/or different time frames ^32^. ***Cluster*** denotes independent isolates from different patients that are within a predetermined SNP threshold of recent common ancestry ^13^ for each NTM species and which warrant further epidemiologic follow-up as potential transmission events ^14^. Isolates that are not monophyletic are considered ***unclustered*.**

### Mycobacterium abscessus subspecies

A total of 203 *M. abscessus* isolates from US CF patients were analyzed, including 155 *M. abscessus* subsp. *abscessus* isolates from 88 patients, 41 *M. abscessus* subsp. *massiliense* isolates from 26 patients and seven *M. abscessus* subsp. *bolletii* isolates from four patients. To compare the sequenced isolates from this study to those from previous reports, we included an additional 15 isolates from 10 separate studies (**Table S2**) in the analysis. Phylogenomic results of the entire sample set, including longitudinal isolates from 41 patients (32 *M. abscessus* subsp. *abscessus*, 7 *M. abscessus* subsp. *massiliense*, and 2 *M. abscessus* subsp. *bolletii*), revealed high sequence similarity among same-patient isolates and clustering on the phylogenetic tree with a mean within-patient diversity of 4.01 SNPs for *M. abscessus* subsp. *abscessus* and 1.62 SNPs for *M. abscessus* subsp. *massiliense*. This is consistent with previous studies that found within-patient isolates of *M. abscessus* subsp. *abscessus* and *M. abscessus* subsp. *massiliense* to be clonal ^13–15,30,31^. Thus, all subsequent analyses were performed using one representative isolate per patient, utilizing the isolate with the maximum read depth and most complete genotypic information. Overall, genetic diversity of *M. abscessus* subsp. *abscessus* among patients ranged from 2-25,537 SNPs out of 3,631,667 core genome positions analyzed, and for *M. abscessus* subsp. *massiliense*, genetic diversity ranged from 2-38,692 SNPs out of 3,960,571 core genome positions analyzed.

Phylogenomic analyses of CF-RDP patient isolates compared to reference isolates revealed the presence of two dominant clones of *M. abscessus* subsp. *abscessus* (**Figure 2A**; MAB clone 1 and MAB clone 2) and two clones of *M. abscessus* subsp. *massiliense* (**Figure 2B**; MMAS clone 1 and MMAS clone 2). As many as 50% (44/88) of patients with *M. abscessus* subsp. *abscessus* had isolates from MAB clone 1. This was the predominant clone observed in a global population study of *M. abscessus* ^14^, a nationwide study of CF-NTM isolates in Italy ^31^, and includes the type strain (ATCC 19977 ^T^) that was isolated from a post-surgical soft tissue infection in 1953 ^33^. MAB clone 2, also identified in European isolate populations (reference strains WRCM9 and UNC673) ^14,30,31^, was observed in 13% (11/88) of US CF patients. The previously described “transmissible” clone of *M. abscessus* subsp. *massiliense* (reference strains 19m, 30a, 08-23-12, 09-19-12 and 2B-0107) ^13,16,17^, called MMAS clone 1, was observed in only 23% (6/26) of patients, while the most predominant strain in 42% (11/26) of patients was MMAS clone 2. MMAS clone 2 has not been described as a dominant clone in any previous studies. None of the CF-RDP *M. abscessus* subsp. *massiliense* isolates were closely related to the type strain CCUG 48898^T^ which was originally isolated from a bronchiectatic patient in France in 2004 ^34,35^. The remainder of CF-RDP *M. abscessus* subsp. *abscessus* (33/88 = 38%) and *M. abscessus* subsp. *massiliense* (9/26 = 35%) patient isolates were unclustered and genetically diverse.

**Figure 2.**
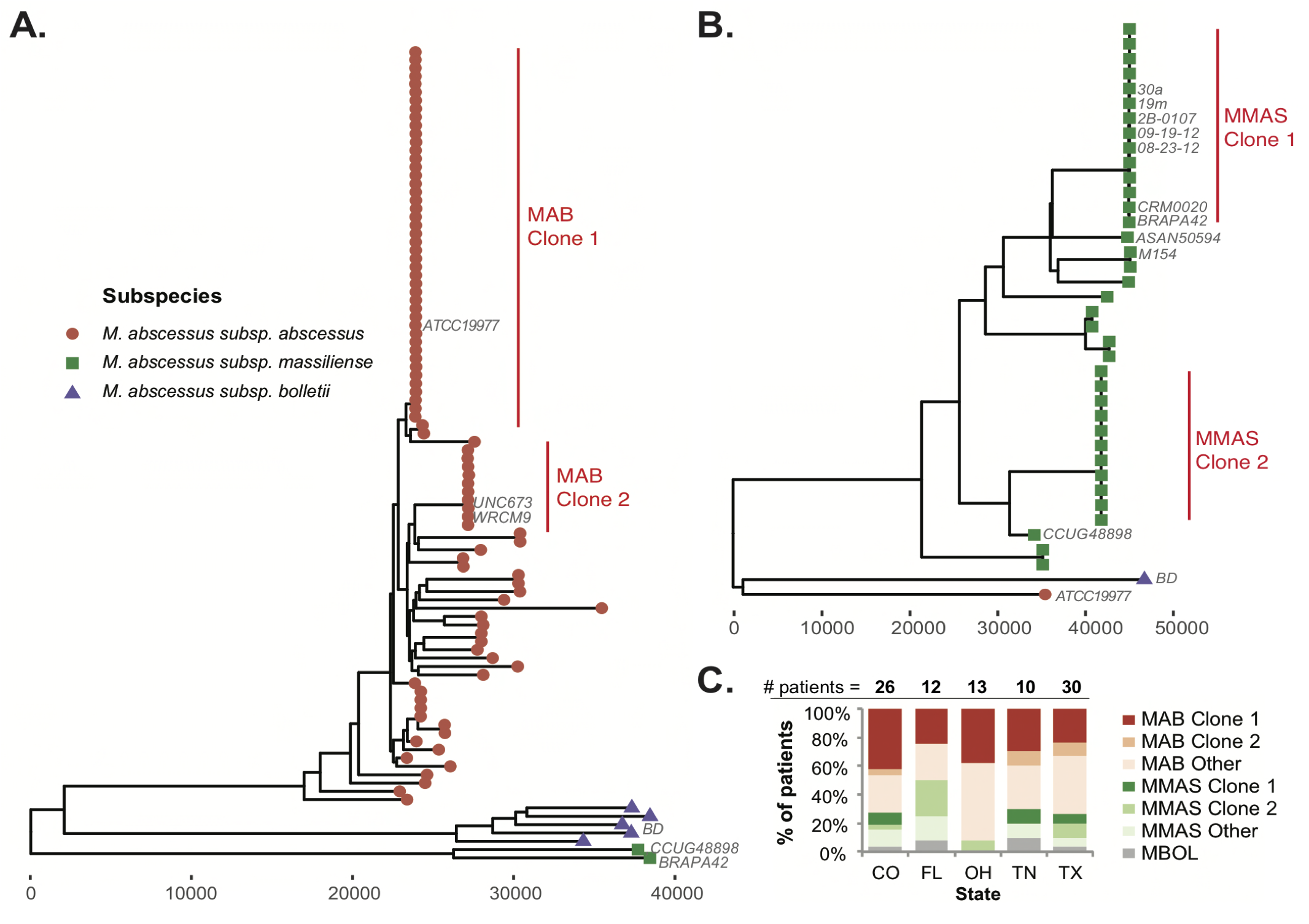
Phylogenomic reconstructions of *M. abscessus* isolates from CF patients using one isolate per patient. Isolates with the highest read depth and most complete genotype information were chosen. A. Core genome phylogenies of *M. abscessus* subsp. *abscessus* isolates and B. *M. abscessus* subsp. *massiliense* with type strains and reference isolates from previous studies (Table S2). Dominant clones are labeled from each group are labeled in red. C. Proportions of clones and unclustered isolate groups from five US states with isolate samples for 10 or more patients. The scale bar represents the number of SNPs.

To determine whether dominant clones or unclustered isolates were enriched in specific geographic areas, we examined the distribution of *M. abscessus* subspecies clones across the entire patient dataset and in states with ten or more patients (**Figure 2C**). Even with limited patient numbers, we observed the *M. abscessus* clones in multiple states spanning different geographic regions, indicating that these clones are widespread across the US. In states with the most patients sampled (CO, FL, OH, TN, TX; **Figure 2C**), we found that MAB clone 1 and MMAS clone 2 were present in all five states and the remaining clones were present in 3-4 states, suggesting a lack of enrichment in particular geographic areas for the dominant clones and unclustered isolates.

### Mycobacterium avium

A total of 80 *M. avium* isolates were analyzed from 46 US CF patients, including 15 patients with longitudinal isolates spanning a range of 2 to 1,052 days from first to last isolate. In contrast to *M. abscessus* subspecies, the mean within-patient diversity of *M. avium* isolates was much higher at 2,216 SNPs. The genetic diversity observed across the entire sample set ranged from 58–17,399 SNPs out of 3,828,484 core genome positions analyzed. A little over half of patients with multiple *M. avium* samples (8/15 = 53%) had clonal isolates with an average within-patient diversity of 9 SNPs, while the remaining patients (7/15 = 47%) cultured two or more strain types over time with an average within-patient diversity of 19,019 SNPs. Among these seven patients, an average of 2.43 strain types per patient was observed (range 2 to 5 strain types).

Phylogenomic analysis of 80 US CF *M. avium* isolates, along with 42 reference and type strains (Table S2), revealed that all US CF isolates were *M. avium* subsp. *hominissuis*, arranged in six monophyletic lineages (Figure 3). No dominant clones of *M. avium* were observed. Of the previously described lineages of *M. avium* that included primarily Asian isolates ^24,25^, only one US CF isolate was observed in the East Asian lineage (reference strain TH135 ^36^). The highest proportion of CF-RDP *M. avium* isolates fell into lineage 3 (30/80 = 37%) followed by lineage 1 (16/80 = 20%), lineage 5 (14/80 = 17%), lineage 4 (12/80 = 15%), lineage 2 (7/80 = 8%), and the East-Asia lineage (1/80 = 1%).

**Figure 3.**
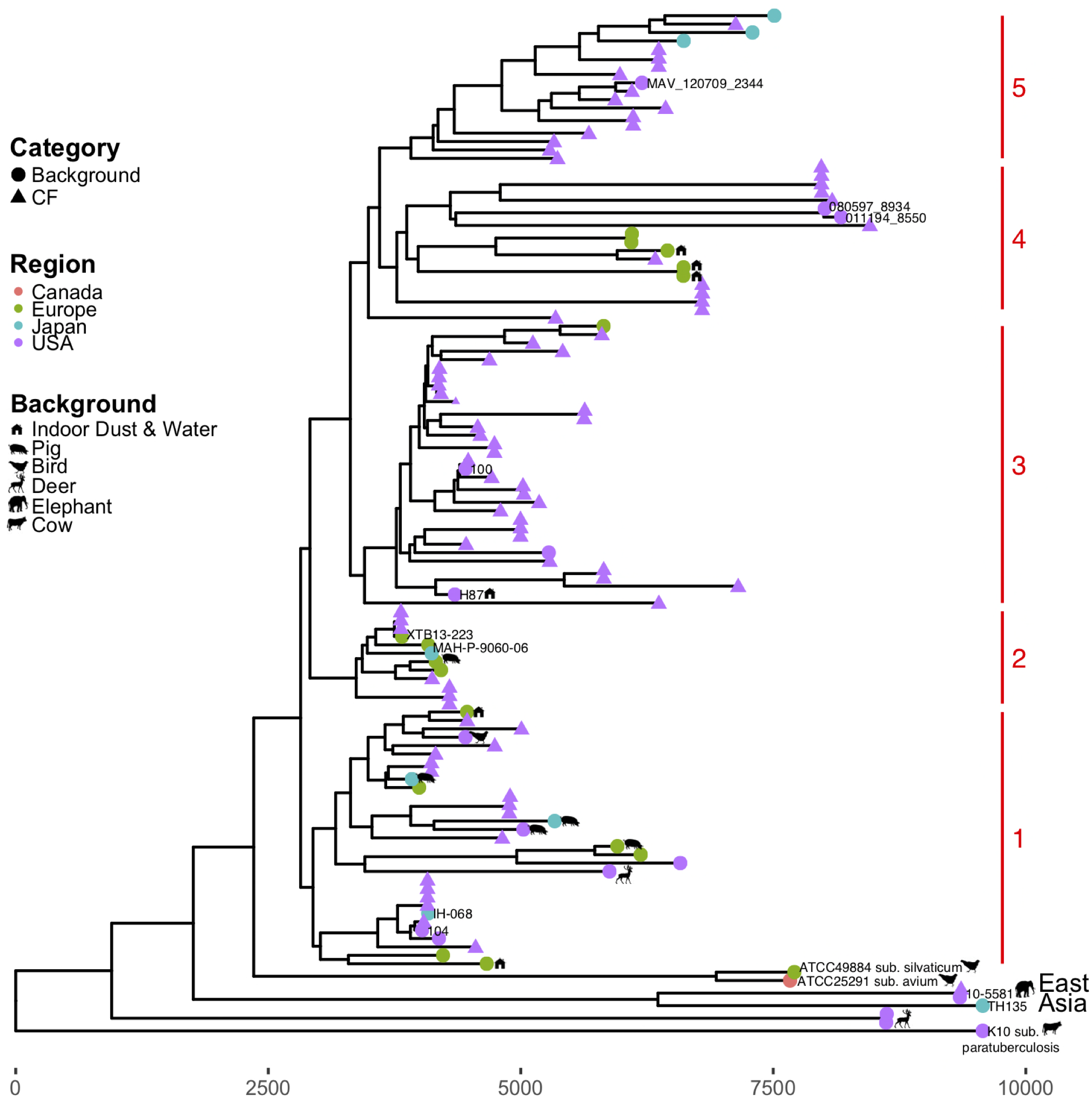
Phylogenomic reconstruction of 80 US CF-RDP *M. avium* isolates and 42 background strains for reference. The six lineages of *M. avium* are labeled on the right. Shapes represent the category of isolate (CF: triangle, Background: circle), the source of background isolate (bird, cow, deer, elephant, pig, indoor dust or water: house) and colors represent the region of origin (Canada: pink, Europe: green, Japan: blue, USA: purple). The scale bar represents the number of SNPs.

To evaluate US *M. avium* isolates and lineages in a global context and among different hosts and reservoirs, we assessed their relationships to genomes from 15 clinical, 6 environmental, and 12 zoonotic isolates from Japan ^24,25,36^, Germany ^26–28^, Belgium ^37^, Belarus (patient; unpublished), Canada (hen; unpublished) and France (pigeon; unpublished; **Table S2**). Lineage 1 included a respiratory isolate (IH-068) from a stable pulmonary patient in Japan ^24^, a virulent isolate collected from a US HIV patient in 1983 (strain 104) ^38^, one isolate from a US deer (unpublished), one isolate from a US bird (unpublished), two isolates collected from two German household dust samples ^28^, four isolates from pigs in Belgium and Japan ^25,37^ and 16 CF-RDP isolates. Lineage 2 included a clinical isolate from Belarus (XTB13-223), a pulmonary isolate from Germany (MAH-P-9060-06), an isolate from a pig in Belgium ^37^, and seven CF-RDP isolates. Lineage 3 contained the highest number of CF-RDP isolates (n=29) along with a laboratory strain with low virulence in mouse models (strain 100) ^38^ and a clinical pulmonary isolate from Germany (strain MAH-P-0913) ^26^. Interestingly, lineage 3 also included strain H87 ^39^, which was isolated from a water source in Colorado and exhibits the ability to infect and survive within free-living amoebae ^40^. Lineage 4 is composed of isolates from patients and dust samples collected in Germany ^26^ and six CF-RDP patient isolates. Lineage 5 includes three pulmonary isolates from Japan ^25^ and 14 isolates from nine US CF patients. The previously described East-Asian lineage had one CF-RDP isolate and a zoonotic isolate collected from an elephant in the US (strain 10-5581; unpublished). In summary, all six lineages included both US and non-US isolates, but lineage 3 had the highest proportion of US isolates. Moreover, lineages 3, 4 and 5 contained no zoonotic isolates, with zoonotic isolates restricted to lineages 1, 2 and East Asia.

### Mycobacterium intracellulare and Mycobacterium chimaera

We analyzed 34 *M. intracellulare* isolates from 24 patients and 12 *M. chimaera* isolates from 11 patients compared to known reference genomes, including *M. chimaera* strains ZURICH1 & 2015-22-71, strains which were implicated in a global outbreak associated with contaminated heater-cooler units (HCUs), and the closely-related species *M. yongonense* (type strain 05-1390^T^; **Figure 4**, **Table S2**). A clear division within the phylogeny distinguishes *M. intracellulare* and *M. chimaera* isolates; however, *M. yongonense* strain 05-1390^T^ phylogenetically places with *M. chimaera* isolates despite a high amount of genetic differences in comparison to CF-RDP *M. chimaera* isolates (mean = 38,288 SNPs) and reference *M. chimaera* isolates (mean = 37,644 SNPs). Only one CF-RDP *M. intracellulare* isolate was genetically-similar (66 SNPs) to a reference isolate (MOTT-02) that was isolated from a 64 year old pulmonary patient in South Korea ^41^. The remaining CF-RDP *M. intracellulare* isolates were related to but genetically distinct from other reference strains (i.e. ATCC 13950 ^42^, MOTT-64 ^43^), zoonotic isolates collected from birds in the US ^44^, and *M. yongonense*. None of the CF-RDP *M. chimaera* isolates were genetically similar to isolates derived from contaminated HCU units mean = 15905 SNPs; ^45,46^. Overall, there seems to be distant relationships between HCU-related, reference isolates and US CF *M. chimaera* and *M. intracellulare* isolates, suggesting these infections come from unrelated sources.

**Figure 4.**
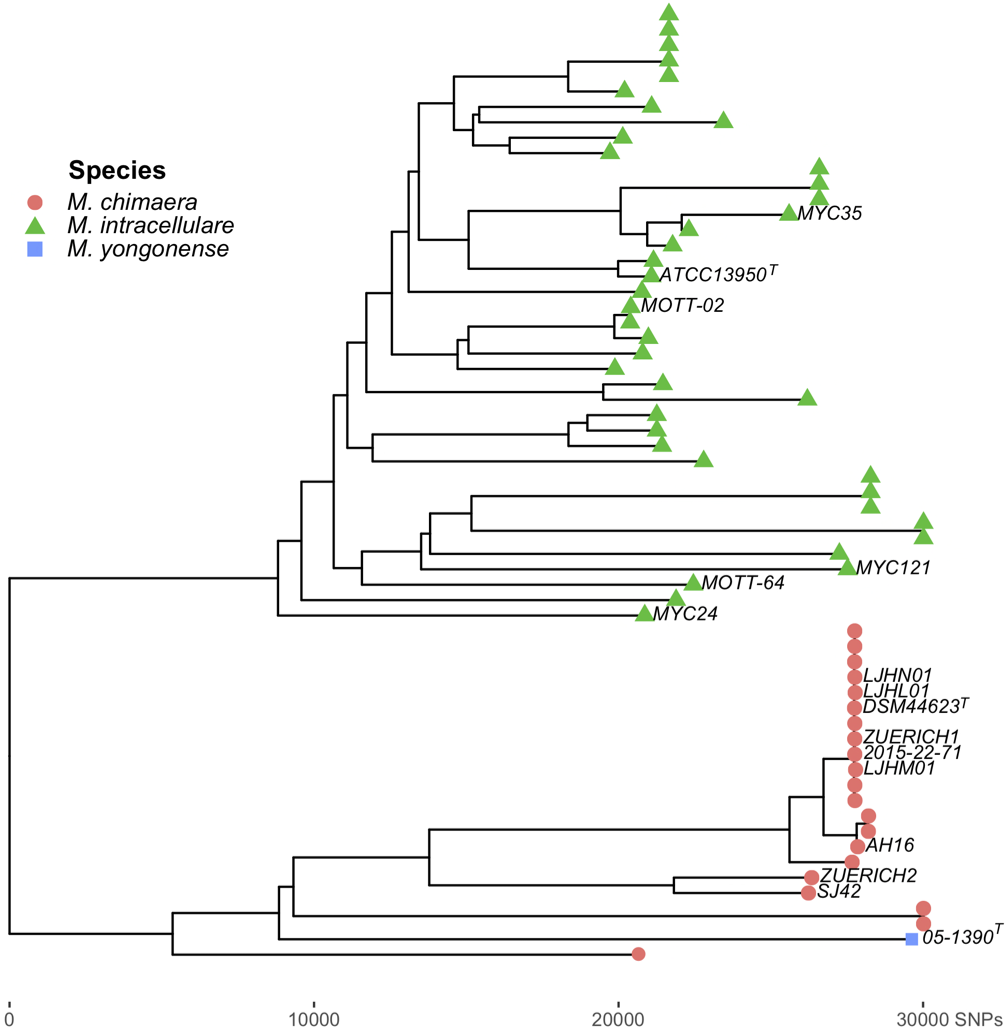
Phylogenomic reconstruction of 34 US CF-RDP *M. intracellulare*, 12 US CF-RDP *M. chimaera*, and 16 background strains. Colored-shapes represent the taxonomic identification of each isolate (*M. chimaera*: red-circle, *M. intracellulare*: green-triangle, *M. yongonense:* blue-square). Phylogenetic relationships are based on 164,693 singlenucleotide polymorphisms (SNPs) identified across genomes compared with the *M. chimaera* CDC 2015-22-71 reference genome. The scale bar represents the number of SNPs.

### Quantitative assessment of strain similarity and potential transmission

In order to identify genetic evidence of potential transmission of NTM between US CF patients and distinguish between widespread genetic “clones”, we examined genome wide SNP distances for all isolate pairs, including longitudinal isolates from the same patients (within-patient) and isolates from different patients (between-patient) for *M. abscessus* subsp. *abscessus, M. abscessus* subsp. *massiliense* and *M. avium*. Comparisons within *M. chimaera* and *M. intracellulare* were excluded from this analysis due to low sample sizes of patients with longitudinal isolates.

Using the pairwise SNP distribution for within-patient isolates as a measure of SNP accumulation or mutation within a known time frame, compared to the SNP distribution of NTM isolates from different patients, we employed a Bayesian statistical approach to calculate the posterior probabilities that patient isolates suggest recent transmission or recent acquisition from a common source. In this calculation, “recent” is defined as the maximum length of the longitudinal samples available (maximum time between collection of patients’ first and last *M. abscessus* subsp. *abscessus* or *M. abscessus* subsp. *massiliense* isolates: 702 days; maximum time between collection of patients’ first and last *M. avium* isolate: 1,052 days). The distribution of within-patient and between-patient NTM genomic SNPs is shown in Figure 4A. For the *M. abscessus* subspecies, we found that between-patient isolates had a 99% probability of recent shared ancestry, suggesting transmission, with a pairwise distance of 5 or fewer SNPs, a 90% probability with 11-15 SNPs and an 80% probability with 16-20 SNPs (**Figure 5**). For *M. avium*, we estimated a 100% probability of recent shared ancestry with 0-15 SNPs, but did not observe any instances of between-patient transmission of *M. avium* at this threshold. The SNP thresholds we propose and evaluate in *M. abscessus* subspecies, are similar to that proposed previously by Bryant et al. ^13^, but our method additionally provides quantitative probability estimates of recent transmission or common acquisition at different SNP thresholds.

**Figure 5.**
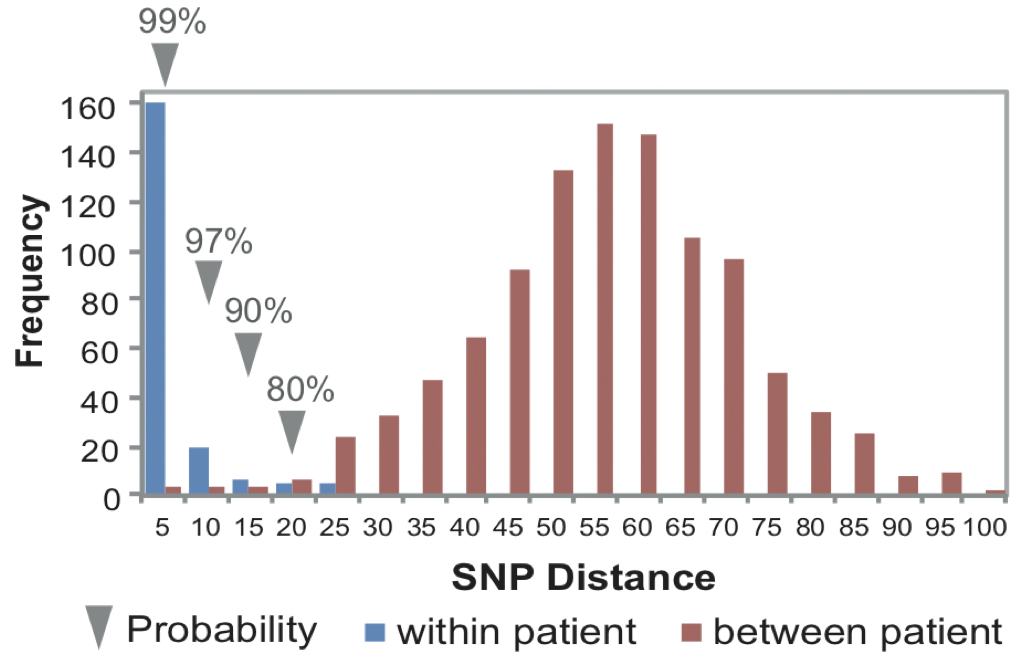
Bayesian cluster analysis of NTM in US CF-RDP patients to identify recent shared ancestry. A. Pairwise SNP distances of *M. abscessus* subsp. *abscessus* and *M. abscessus* subsp. *massiliense* isolates from within same patients (blue) and between different patients (red) are shown. Estimated probabilities of recent shared ancestry (compared to within-patient isolates with known time frames) are labeled with grey arrows.

Using the 80% probability threshold for recent shared ancestry, six clusters of patients with genetically similar *M. abscessus subsp. abscessus* isolates and four clusters of patients with genetically similar *M. abscessus* subsp. *massiliense* isolates were identified for a total of 10 genetically similar clusters. By subspecies, 16/88 patients with *M. abscessus* subsp. *abscessus* (18%) and 10/26 patients with *M. abscessus* subsp. *massiliense* (38%) belonged to clusters within the threshold of recent shared ancestry and warranted epidemiological follow-up. Only 12 of the patients in 5/10 *M. abscessus* clusters were seen in the same facility, including 3 facilities in 2 states (CO, TX). Overall, only 12/114 patients (10.5%) with *M. abscessus* subspecies in the study had isolates that were genetically similar and seen at the same facility. In contrast, no genetically similar *M. avium* isolates were identified between US CF patients as the within and between-patient distributions of SNP distances did not overlap and the lowest SNP distance between-patients was 58 SNPs.

## Discussion

To address concerns of potential transmission of NTM between CF patients and identify instances of genetically similar isolates, the Colorado CF-RDP was initiated by the CF Foundation in 2015 as a voluntary nationwide surveillance program for CF centers. During the first two years of surveillance, WGS was performed on 341 NTM isolates sent from 45 CF centers in 22 states across the US. Analysis of this isolate cohort provides an expanded view of the genomic diversity and population structure of the CF-NTM population within the US, including the most predominant NTM species, *M. abscessus* and *M. avium*, and the less common taxa, *M. abscessus* subsp. *massiliense, M. intracellulare* and *M. chimaera*. In addition, these data allow us to examine whether genomic trends from previous European, Australian, and Asian studies are mirrored within US NTM populations.

Recently, it was shown that two dominant clones of *M. abscessus* and a potentially human-transmissible clone of *M. abscessus* subsp. *massiliense* ^13,17,29^ make up a substantial portion of the clinical isolate population in Europe, Australia, and a single location in the US (North Carolina) ^14^. Our results show that a majority of US CF *M. abscessus* isolates are members of four dominant clones, including a newly identified clone of *M. abscessus* subsp. *massiliense*, MMAS clone 2. To refine the method for identifying genetic evidence of potential transmission beyond monophyletic assignments of clones within a species, we developed a Bayesian statistical method to calculate the posterior probability of recent shared ancestry between patient isolates, indicative of shared source acquisition or transmission between patients. Using this method, we identified 10 clusters involving 26 patients with 80% probability of recent shared ancestry in the Colorado CF-RDP study cohort. The epidemiologic data, limited to collection dates and facility locations, shows that less than half of the clusters include 12 patients that were seen in the same CF centers. That only 10.5% of patients with *M. abscessus* subspecies (12/114 patients) had both genetic and limited epidemiological evidence for potential transmission suggests that patients are not likely to be acquiring NTM at a high rate during CF care, as was also observed in a nationwide surveillance study in Italy ^31^. Furthermore, the genetically similar isolates in our study are geographically diverse and may simply be acquired through common geographic or environmental locations, though we did not have access to all residential locations and potential for social contact among CF subjects to test these hypotheses.

The mechanism of spread of highly similar *M. abscessus* isolates within the US remains unclear. Hypotheses for the observed similarity between CF patients have been proposed including direct transmission between CF patients in the clinic or indirect transmission through a shared contaminated space, fomite spread, or the generation of long-lived infectious aerosols ^14^. A currently unexplored hypothesis is a systematic study of the geographic diversity and exposure of these clones in the environment. Indeed, the literature is scant with examples of *M. abscessus* found in the environment, with the exception of a handful of studies in Australia ^47,48^, Hawaii ^49^, and New York ^50^ where *M. abscessus* was isolated from plumbing biofilms and water samples. The lack of matched environmental NTM surveillance and detailed epidemiological information in the current study limits our ability to empirically test the hypothesis of patients independently acquiring highly similar isolates from a widespread genotype in the environment versus nosocomial transmission. In either case, these clones represent strains that are geographically widespread, genetically stable and evolutionarily adapted for human infection.

This study also provides the first report of genomic epidemiology for *M. avium* and other MAC species in US CF patients. In comparison to the population structure and highly similar clones of *M. abscessus* in US CF patients, *M. avium, M. intracellulare and M. chimaera* isolates were genetically diverse and largely unclustered. Additionally, US *M. avium* isolates were, by and large, distinct from clinical populations and lineages described previously in Asian studies ^24,25^. US lineages of *M. avium* also grouped with reference isolates from non-CF patients, environmental samples from Europe ^27,28^, and zoonotic isolates from Belgium ^37^, Canada, France and Japan ^25^ suggesting that similar clades have been observed in other parts of the world among diverse hosts. However, the majority of *M. avium* observed in US CF patients are genetically dissimilar enough (>600 SNPs) from the zoonotic isolates included in this study suggesting that patients and animals in the US may be acquiring their NTM infections from different reservoirs. In contrast with *M. abscessus*, in which within-patient isolates are clonal over time, nearly half of patients with *M. avium* cultured multiple strains over time. Taken together, we surmise that *M. avium* isolates found in US CF patients are likely not derived from patient-to-patient transmission, but instead from independent acquisition of genetically diverse strains in the environment; our data corroborate previous work showing genetically matched environmental and patient *M. avium* isolates ^51–57^. Two hypotheses can explain the observations of multiple genotypes and species in CF patients: 1) patients are originally infected with multiple genotypes of MAC that are selected for during infection and treatment, or 2) patients clear the original infection and subsequently acquire a new, independent genotype from environmental exposure. Further studies of within-patient population diversity with corresponding environmental sampling are needed to address these questions.

The taxonomy within MAC continues to evolve. A recent publication demoted *M. yongonense* to *M. intracellulare* subsp. *yongonense* ^58^, but did not compare *M. yongonense* 05-1390^T^ to any *M. chimaera* genomes. Within our MAC isolate comparisons, we found *M. yongonense* 05-1390^T^ phylogenetically placed within the *M. chimaera* clade. Previous research suggests that *M. yongonense* has distinct genotypes ^59^ and lateral gene transfer of loci e.g. rpoB; ^60^, which may explain the unexpected phylogenetic placement of *M. yongonense* 05-1390^T^ among our *M. chimaera* genomes. Further examination of MAC isolates, with specific attention to the relationships between *M. chimaera, M. intracellulare* and *M. yongonense*, will clarify the taxonomy of these species and allow us to better assess their prevalence within patients.

Our research demonstrates the utility of WGS to perform NTM surveillance, uncover potential instances of transmission between patients, and assess the dynamics of NTM infections in CF patients. The findings of our initial surveillance work of NTM in US CF patients underscore the need for continued epidemiologic follow-up in patients with *M. abscessus* subsp. *abscessus* and *M. abscessus* subsp. *massiliense* to assist infection control measures and limit the spread of NTM infections where possible. The different epidemiologic patterns observed between *M. abscessus* and MAC species suggest that different interventions may be required to limit patient exposure to these emerging pathogens.

## Materials and Methods

### Bacterial samples

NTM isolates from CF patients in the United States were sent to the US CF-RDP NTM Culture and Biorepository Core within the Mycobacteriology Reference Lab at National Jewish Health for processing. All samples were cultured on solid media prior to sub-culturing single colony isolates that were divided into eight 1mL biobanked glycerol stock aliquot replicates stored at −20 Celsius in the Culture and Biorepository Core of the CF-RDP. DNA was isolated from all isolates for whole genome sequencing.

### DNA Extraction and Whole-Genome Sequencing

NTM DNA was isolated using a modified protocol from Käser et al. ^61^, replacing the phenol:chloroform extraction with a DNA column cleaning approach (Genomic DNA Clean & Concentrator, Zymo Research, Irvine, CA). Illumina sequencing was performed by the Colorado CF-RDP NTM Molecular Core at National Jewish Health. The NexteraXT DNA sample preparation kit was used to prepare genomic DNA (gDNA) libraries (Illumina) and quality was assessed via examining the distribution of fragment lengths using BioAnalyzer. Genomic Illumina libraries were multiplexed for sequencing using MiSeq v3 sequencing and 2×300 read lengths.

### Single Nucleotide Polymorphism (SNP) Analysis

llumina reads were trimmed of adapters and base calls with quality scores less than Q20 using Skewer ^62^. Trimmed reads were assembled into scaffolds using Unicycler ^63^. Genome assemblies were compared against a selection of reference genomes to estimate average nucleotide identities (ANI) and assign a species call to each isolate ^64,65^. A cutoff ANI of ≥ 95% indicated the isolate and reference genome belonged to the same species.

Trimmed sequence reads were mapped to respective reference genomes and compared as previously described ^15,66^. Isolates identified as *M. abscessus* subsp. *abscessus* and *M. abscessus* subsp. *bolletii* were mapped to *M. abscessus* reference genome of strain ATCC 19977^T 33^; all isolates identified as *M. abscessus* subsp. *massiliense* were mapped to complete genome of *M. abscessus* subsp. *massiliense* strain BRA_PA_42 ^67^; all isolates identified as *M. avium* were mapped to the genome of *M. avium* strain H87 ^39^; and all isolates identified as *M. chimaera* or *M. intracellulare* were mapped to the genome of *M. chimaera* CDC 2015-22-71 ^45^.

For each taxon (e.g. *M. avium*), background reference genomes are also included by *in silico* creation of sequence reads of the background genome sequences available on NCBI (Table S2). *In silico* and Illumina reads were then mapped to the designated reference genome using Bowtie2 software ^68^. SNPs were called using SAMtools mpileup program ^69^ and were filtered using a custom perl script using the following parameters: SNP quality score of 20, minimum of 4x read depth, the majority of base calls support the variant base, and less than 25% of variant calls occur at the beginning or end of the sequence reads.

A multi-fasta sequence alignment was created from concatenated base calls from all isolates. SNP-sites was used to filter positions in the genome in which at least one strain differed from the reference genome, and for which high quality variant and/or reference calls were present for all other strains ^70^.

### Phylogenetic Analysis

Concatenated sequences were used to make phylogenetic trees using Maximum-Likelihood with 500 bootstrap replicates in RAxML-NG using nucleotide general time-reversible model; ^71,72^. Phylogenetic trees were annotated and visualized with ggtree ^73^. Non-CF reference strains were included to allow for comparison to previous studies. For patients with multiple *M. abscessus* isolates sequenced, the isolate fasta sequence with the highest read depth and least amount of missing data (insertion/deletions or N’s) was selected and used for subsequent one isolate per patient comparisons.

### Bayesian Analysis for Recent Shared Ancestry

For each taxon, pairwise SNP distances were calculated between the core genomes of isolates collected from US CF patients using MEGA ^74^. Each pair within each species was classified as from longitudinally isolates collected from a single patient (within patient) or isolates collected from different patients (between patients). Frequencies of pairwise SNP distances between longitudinally sampled isolates within patients were calculated, binned in 5 SNP increments and used to calculate a Bayesian posterior probability that two between-patient strains are of recent common ancestry (H = Hidden) suggestive of transmission or common acquisition, based on the Observed (O) SNP differences between strains, using the following equation:

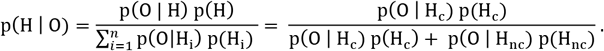

The distributions of the pairwise genomic SNPs from both the within-patient and between-patient pairwise comparisons (Figure 4) are used to calculate the likelihoods p(O | H), defined as the probability of the Observed (O) given the Hidden (H), where the Observed is the genomic SNP difference between NTM pairs, and the Hidden is that they are either clonal (Hc) originating from the same patient, or not clonal (Hnc) originating from different patients. A uniform prior p(H) of 0.5 was used in this equation, since we do not know *a priori* the frequency of NTM transmission events occurring. This posterior probability estimates an empirical threshold used to identify pairs of genetically similar isolates shared between two or more patients that warrant follow-up epidemiological investigation. Analogous Bayesian inference methods, using genomic evidence, have been used previously to identify infectious disease transmission chains ^75^.

Ethical approval for this work was obtained from the National Jewish Health Institutional Review Board through HS-3149.

WGS data are available at the National Center for Biotechnology Information (NCBI) under BioProject PRJNA319839. The Colorado CF-RDP is a research resource for NTM isolates and WGS data for the CF community. More information about the Colorado CF-RDP and the NTM isolates described in this study can be found at https://www.nationaljewish.org/cocfrdp.

## Acknowledgements

This work was supported by the US Cystic Fibrosis Foundation Colorado Research Development Program Grant (NICK15RO) at National Jewish Health. RMD was funded by NIH-NIAID K01-AI125726. The funders had no role in study design, data collection and analysis, decision to publish, or preparation of the manuscript.

## Supplementary Tables

**Table S1**. Metadata for CF-RDP isolates sequenced in the study cohort. Attributes for all the analyzed *M. abscessus, M. avium, M. chimaera* and *M. intracellulare* genomes include isolate identifier, source, country and region of origin, the clade/lineage, Bayesian-defined cluster, reservoir, taxonomic identification, and SRA number.

**Table S2**. Metadata for reference, type strain and background isolates used in this study. Attributes for all the analyzed *M. abscessus, M. avium, M. chimaera* and *M. intracellulare* genomes include isolate identifier, source, country and region of origin, clade/lineage, Bayesian-defined cluster, reservoir, taxonomic identification, and citation.

